# Depressive disorder and its associated factors among prisoners in Debre Berhan Town, North Showa, Ethiopia

**DOI:** 10.1101/703223

**Authors:** Yared Reta, Ruth Getachew, Melese Bahiru, Bethelhem Kale, Keralem Workie, Yohannes G/Egziabher

**Affiliations:** Department of Psychiatry, Hawassa University, Hawassa, Ethiopia; Arada Sub-city, HIV prevention, and control office, Addis Ababa, Ethiopia; Addis Ketema Sub-city, Health center, Addis Ababa, Ethiopia; Ginella Health center, Ginella woreda, Harari, Ethiopia; School of Nursing, Debre Berhan University, Debre Berhan, Ethiopia

**Author notes:** Corresponding author: YR. **Email addresses**: RG, MB, BK, KW, YG. These authors contributed equally to this work (Ruth Getachew, Melese Bahiru, Bethelhem Kale, Keralem Workie, Yohannes G/Egziabher).

**Keywords:** Depression, Prison, Inmates, North Showa, Ethiopia

## Abstract

**Background:** Depression is a commonest mental disorder among prisoners characterized by an intense mood involving a feeling of sadness, lack of interest or hopelessness that lasts for weeks, months, or even longer. In addition to imprisonment, depression is the primary factor leading to suicidal attempt. Therefore, this study revealed the magnitude of depressive disorder and its associated factors among prisoners of Debre Berhan Town.

**Methods:** We conducted an institution based cross-sectional quantitative study. We collected the data from 336 randomly selected prisoners by using interviewer-administered Patient Health Questioner-9 (PHQ-9). Multiple logistic regression was performed to identify independent predictors.

**Result:** Out of the total of 336 prisoners 330 (98%) were males. Using PHQ-9 at the cut of point >5 for caseness, the prevalence of depression found to be 44% (n=148). Widowed (AOR=6.30 CI: 1.09-36.67), those who are educated at college or university level (AOR=5.34 CI:1.59-17.94), a history of suicidal attempt (AOR=2.76 CI: 1.04-7.31), Previously facing severe stressful life event (AOR=2.57 CI: 1.41-4.67), 5-10 years of sentence (AOR=2.51 CI:1.32-4.79) and having chronic medical illness (AOR= 3.32 CI: 1.26-8.75) are found to be independently associated with depression.

**Conclusion:** In general, there is a high prevalence of depression among prisoners in of Debre Berhan town. Therefore, designing strategies for early screening and treatment of depression at prisons is very crucial.

## Introduction

More than 10 million people are imprisoned worldwide. The prevalence of a mental disorder among prisoners is at least five times the rate in the general population, that the clinical staff ignores a large proportion of inmates with severe mental disorders, and that those who do receive treatment do not receive adequate attention [1–3]. Once imprisoned, prisoners are no longer free to choose living places and how to spend their time. Deprivation of liberty as a result of imprisonment invariably results in denial of choices usually taken for granted in the outside community [4, 5].

There are external and internal factors that exacerbate mental illness morbidity in prisons. Common external factors are overcrowding and unhygienic living conditions, poor quality of food, inadequate health care, physical or verbal aggression by inmates, availability of illicit drugs, lack of privacy and time for quiet relaxation. Internal factors are mostly emotional, where prisoners may have feelings of guilt or shame about the offenses they have committed, experience stigma of being imprisoned, worry about the impact of their behavior on others, including their families and friends. The cumulative effect of all these factors left the prisoner unchecked, tends to worsen their mental health and increases the likelihood of damage to the wellbeing of the prisoners [1, 4].

Depressive disorders are a significant contributor of suicide characterized by a feeling of sadness, loss of interest or pleasure, feelings of low self-esteem or guilt, disturbance of sleep or appetite, decreased energy, and poor concentration. Depression can be chronic or short lasting, markedly impairing an individual’s functioning at work or school or cope with daily life. In 2015 World Health Organization (WHO) estimated that 4.4% (322 million people) of the global population live with depression. [6, 7].

Suicide is the 10th leading cause of death in the US (more common than homicide) and the third leading cause of death for ages 15 to 24 years [8]. Incarcerated individuals are more likely to attempt and commit suicide than individuals living in the general population [9]. Suicide is often the single most common cause of death in prisons, and more than 90% of those who die by suicide had one or more mental disorders with depression the commonest one [10, 11]. A systematic review from 12 countries s revealed that prisoners were many time more likely to have a depressive disorder than the general population [12].

A study from Iran, among 351 inmates, 29% met Diagnostic and Statically Manual Four Text Revision (DSM-IV TR) criteria for a current diagnosis of MDD [13]. In Brazil, using the Mini International Neuropsychiatric Interview(MINSI) in the sample of 497 prisoners, the prevalence rate of depression found in the closed and semi-open prison systems were 17.6% and 18.8%, respectively [14]. In Eastern Nepal, a cross-sectional study revealed the prevalence of depression among 434 randomly selected inmates was 35.3% [15].

A study done in Durban, South Africa, among 193 prisoners using the MINSI, lifetime prevalence of depression was 24.9 % [16]. A study in an Egyptian prison; using Psychiatric Symptoms Checklist-90 (SCL-90) revealed a high prevalence of depression 82.5% [17]. A study in Nigeria maximum security prison showed the prevalence of psychiatric disorder to be 57% and depression accounts for 30.8% [18].

A cross-sectional study (n=355) in Hawassa Central Correctional Institution determined the prevalence of depression among prisoners by using the Patient Health Questionnaire (PHQ-9) is 56.4% [19]. Another most recent study from Ethiopia (Bahir Dar Prison) using the same tool (PHQ-9) revealed the prevalence of depression 45.5% [20] and a similar prevalence of depression is observed in Jimma Town Prison (41.9%) [21] and the prisons of Northwest Amhara, Ethiopia (43.8%) [22].

Different risk factors may result in depression such as gender and age; economic, educational, employment and marital status; disability, and poor social support; chronic medical illness (like, hypertension, epilepsy, HIV/AIDS); history of suicide, family history of psychiatric problems and exposure to violence and crime, and acculturation stress [23–31].

Despite the existence of many prisons in Ethiopia, to the best of the authors understanding, there is no published research done in the field before the study period. Depression causes much suffering as well as long-term adverse consequences if left untreated. Sadly, depression is hardly ever recognized and managed in prison settings [32]. Therefore, to expose the ground fact, we studied the prevalence of depression and associated factors among Debre Berhan prisoners.

## Methods and Materials

### Study design and Setting

We used an institutional based cross-sectional study design. The Data was collected in May 2015 from Debre Berhan prison, North Showa, Amhara Region, Ethiopia. Debre Berhan town, the capital of North Showa zone, is located in the North Showa zone of Amhara regional state. It is 695 km far from the capital of Amhara Regional State (Bahirdar) and 130 km from the capital of Ethiopia (Addis Ababa). Debre Berhan town has one prison which was established in 1948 G.C. During the period of data collection there were about 1,738 (1704 males and 34 females) inmates in prison.

### Participants and sample size

We used a single population proportion formula to calculate the sample size with an assumption of the prevalence of depression, 50%, which yield adequate sample size. We took 5% as the minimal difference with 95% confidence of certainty. Since our source population is 1,738, we used adjustment formula. Thus, the calculated sample size was 314 and adding a 10% non-response rate, and the total sample size became 345. We selected study participants randomly (using OpenEpi Random Number Generator) by taking prisoners list from prisoners’ registration book.

### Data collection procedure

#### Study Variables

A depressive disorder is the outcome variable. The exposure variables included: Socio-demographic and economic factors such as age, sex, marital status, ethnicity, religion, average monthly income, educational status, occupation and social support (poor support, loneliness). Psychological factors include childhood abuse, family history of mental illness, history of self-harm, and stress. Imprisonment status includes: being imprisoned previously, duration of imprisonment, frequency of imprisonment, type of sentence, type of crime, type of confinement (close custody).

### Measurements

We collected the data using a structured interviewer-administered the questionnaire. We used the Patient Health Questionnaire (PHQ-9) to assess the outcome variable, depressive disorder. PHQ-9 is a structured questionnaire which measures depressive disorder, extracting 9 of the major depressive disorder symptoms from DSM-IV criteria. PHQ-9 sensitivity of 88% and a specificity of 88% for major depression [3, 33]. PHQ-9 is also found to be a reliable and valid tool in different countries, including Ethiopia [34–37]. PHQ-9 consists of 9 items, and each item response is rated as “0” (not at all) to “3” (nearly every day), and the score can range from 0 to 27 [3, 36]. And Social support was determined by using Oslo-3 item social support scale with items sum > 9 showing good social support [38].

To ensure completeness and consistency inflow of items, to minimize systematic errors and to estimate the time needed to complete the questioner; we pre-tested the questioner on 5% of a similar population in another department before the actual data collection.

### Statistical analysis

We checked the data completeness and entered to EpiData (Classic) Entry version 3.1 and exported the data to IBM SPSS version 24 for cleaning and further analysis. We used percentage and frequencies to present descriptive statistics. Variables associated at binary logistic regression (P<0.05), were computed to multiple logistic regression to control confounders. We used the adjusted Odds ratio to measure the strength of association between explanatory and outcome variables at a significance level <0.05.

### Ethical statement

Ethical clearance was obtained from Hawassa University College of Medicine and Health Sciences Institutional Review Board. We have got written consent and signature from respondents before starting the questionnaire.

## Result

### Socio-demographic characteristics of the prisoners

Out of 345 participants, we got complete data found from 97.39 % (n=336). 98.2% (n=330) of the participants were males. Nearly 80% of the sample are age less than 34 (n= 268) and majorities 59.2% (n=199) are single. 94.9% (n=319) are Amhara by ethnicity, and 92% (n=312) of the respondents are Ethiopian orthodox followers. The large proportion 74.7% (n=251) of the prisoners were employees before imprisonment; the majority 72.9 % (n=245) had monthly income of less than 500 Ethiopian Birr and about 57.4 % (n=193) had attended primary school. (Table 1)

**Table 1:** Socio-demographic characteristics of inmates of Debre Berhan prison, North Showa, Amhara Region, Ethiopia, 2015.

| Variables |  | Frequency | % |
| --- | --- | --- | --- |
| Sex | Male | 330 | 98.2% |
|  | Female | 6 | 1.8% |
| Age | 18-24 Years | 134 | 39.9% |
|  | 25-34 Years | 134 | 39.9% |
|  | 35-44 Years | 50 | 14.9% |
|  | =>45 Years | 18 | 5.4% |
| Marital status | Single | 199 | 59.2% |
|  | Married | 115 | 34.2% |
|  | Widowed | 10 | 3.0% |
|  | Divorced | 12 | 3.6% |
| Religion | Orthodox | 312 | 92.9% |
|  | Protestant | 4 | 1.2% |
|  | Muslim | 20 | 6.0% |
| Ethnicity | Amhara | 319 | 94.9% |
|  | Oromo | 13 | 3.9% |
|  | Tigray | 4 | 1.2% |
| Educational status | Illiterate | 52 | 15.5% |
|  | Primary school | 193 | 57.4% |
|  | Secondary | 69 | 20.5% |
|  | College or University | 22 | 6.5% |
| Occupation | Student | 68 | 20.2% |
|  | Unemployed | 85 | 25.3% |
|  | Farmer | 64 | 19.0% |
|  | Government employed | 26 | 7.7% |
|  | NGO | 50 | 14.9% |
|  | Daily laborer | 31 | 9.2% |
|  | Other (Retired and housewife) | 12 | 3.6% |
| Average monthly income | <500birr | 245 | 72.9% |
|  | 500-1000birr | 45 | 13.4% |
|  | 1001-1500birr | 11 | 3.3% |
|  | 1501-2000birr | 12 | 3.6% |
|  | >2000birr | 23 | 6.8% |

### Psycho-social, imprisonment and physical health status of the prisoners

17% (n=57) of the prisoners responded yes for a history of childhood abuse, and 6% (n= 20) responded yes for the family history of mental illness. 8.9% (n=30) had at least a history of single suicidal attempt, and around one third 28%(n=94) of the participants has passed through a stressful life event. This is their second or more imprisonment for 5.4% (n=18) of the respondent. Homicide 47.6% (160) and stealing and robbery 33.0% (111) were found to be the two leading reasons for imprisonment. (Table 2)

**Table 2:** Psycho-social, imprisonment and physical status of inmates of Debre Berhan prison, North Showa, Amhara Region, Ethiopia, 2015.

| Variables |  | Frequency | % |
| --- | --- | --- | --- |
| History of childhood abuse | Yes | 57 | 17.0% |
|  | No | 279 | 83.0% |
| Family history of mental illness | Yes | 20 | 6.0% |
|  | No | 316 | 94.0% |
| History of suicidal attempt | Yes | 30 | 8.9% |
|  | No | 306 | 91.1% |
| Ever had stressful life event | Yes | 94 | 28.0% |
|  | No | 242 | 72.0% |
| Type of stress full life event | Death of beloved one | 40 | 42.6% |
|  | Being in prison | 22 | 23.4% |
|  | Money or material loss | 32 | 34.0% |
| Past history of imprisonment | Yes | 18 | 5.4% |
|  | No | 318 | 94.6% |
| Duration of current sentence | <5 Years | 109 | 32.4% |
|  | 5-10 Years | 99 | 29.5% |
|  | >10 Years | 128 | 38.1% |
| Duration of current imprisonment | <5 Years | 264 | 78.6% |
|  | 5-10 Years | 72 | 21.4% |
|  | >10 Years | 0 | 0.0% |
| Reason for current imprisonment | Homicide | 160 | 47.6% |
|  | Stealing and robbery | 111 | 33.0% |
|  | Rape | 31 | 9.2% |
|  | Abduction | 19 | 5.7% |
|  | Corruption | 15 | 4.5% |
| Physical disability | Yes | 30 | 8.9% |
|  | No | 306 | 91.1% |
| Types of disability | Hearing or Visual | 10 | 33.3% |
|  | Walking | 20 | 66.7% |
| History headaches during the past one month | Yes | 57 | 17.0% |
|  | No | 279 | 83.0% |
| History back pain during the past one month | Yes | 43 | 12.8% |
|  | No | 293 | 87.2% |
| History of fever during the past one month | Yes | 48 | 14.3% |
|  | No | 288 | 85.7% |
| Other health problems (HPTN, DM, Epilepsy) | Yes | 35 | 10.4% |
|  | No | 301 | 89.6% |

From the study participants, 8.9 % (n=30) have a physical disability. Those who often have headache, back pain, and fever, over the past 30 days constitute 17 % (n=57), 12.8 % (n=43), & 14.3 % (n=48) of the study participants respectively. According to Oslo 3-item social support scale those who have poor, moderate, & strong social support constitute 79.2 % (n=266), 19.9 % (n=67) & 9 % (n=3), respectively. (Table 2)

### Prevalence of depression among prisoners

Using a cut-off point of ≥ 5 for detecting cases on PHQ-9, the point prevalence of depression among prisoners of Debre Berhan town prison was found to be 44% (n=148) (95% CL: 38.8%, 49.9%).

### Determinants of depression among prisoners of Debre Berhan town prison

During Bivariate analysis of socio-demographic variables marital status, educational status & religion and income before imprisonment were significantly associated with depression. Bivariate analysis of physical health status and social supports shows that those who had a headache, back pain, and fever over the past 30 days and those who have any chronic health problem were found to have a significant association with depression.

Multiple logistic regressions were used to minimize the risk of confounder and variables with P-value of < 0.05 are reported to be significantly associated with depression. The odds of having depression is found to be higher in widowed than in singles (AOR=6.30 CI: 1.09-36.67). Those prisoners educated at the college level or university level are more likely to have depression than illiterates (AOR= 5.34 CI: 1.59-17.94). (Table 3)

**Table 3:** Simple and multiple binary logistic regression analysis showing significant predictors of of inmates of Debre Berhan prison, North Showa, Amhara Region, Ethiopia, 2015.

| Variables | Category | Depression |  | OR with 95% CI |  |
| --- | --- | --- | --- | --- | --- |
|  |  | No | Yes | Crude | Adjusted |
| Age | 18-24 Years | 70 | 64 | 0.35 (0.12-1.04) | 1.68(0.42- 6.73) |
|  | 25-34 Years | 87 | 47 | 0.21 (0.07-0.62) * | 0.86(0.23-3.24) |
|  | 35-44 Years | 26 | 24 | 0.36 (0.11-1.15) | 1.26(0.30-5.22) |
|  | ≥45 Years | 5 | 13 | 1 | 1 |
| Marital status | Single | 117 | 82 | 1 | 1 |
|  | Married | 64 | 51 | 1.26 (0.71 – 2.24) | 1.43(0.80-2.57) |
|  | Widowed | 2 | 8 | 5.80(1.02 – 33.87) * | 6.30(1.09-36.67) * |
|  | Divorced | 5 | 7 | 2.25(0.56 - 8.94) | 1.88(0.44-7.95) |
| Educational status | Illiterates | 33 | 19 | 1 | 1 |
|  | Primary school | 114 | 79 | 1.20(0.64 - 2.27) | 1.05(0.52-2.15) |
|  | Secondary School | 35 | 34 | 1.69(0.81 - 3.52) | 1.32(0.57-3.12) |
|  | College/ University | 6 | 16 | 4.63 (1.55-13.84) * | 5.34 (1.59-17.94) * |
| Ever sustained childhood abuse | Yes | 25 | 32 | 1.8 (1.01 - 3.20) * | 1.34(0.66-2.76) |
|  | No | 163 | 116 | 1 | 1 |
| History of suicidal attempt | Yes | 8 | 22 | 3.93 (1.70 - 9.12) ** | 2.76(1.04-7.31) * |
|  | No | 180 | 126 | 1 | 1 |
| Ever had serious stressful life event | Yes | 31 | 63 | 3.75(2.267 - 6.217) ** | 2.57(1.41-4.67) * |
|  | No | 157 | 85 | 1 | 1 |
| Previously imprisoned | Yes | 5 | 13 | 3.52 (1.23 - 10.12) * | 2.94(0.84-10.29) |
|  | No | 183 | 135 | 1 | 1 |
| Duration of current sentence | <5 years | 71 | 38 | 1 | 1 |
|  | 5-10 years | 46 | 53 | 2.15 (1.23 - 3.76) * | 2.51(1.32-4.79) * |
|  | > 10 years | 71 | 57 | 1.50 (0.89 - 2.54) * | 1.04(0.55-1.95) |
| Headache over the past 30 days | Yes | 21 | 36 | 2.556 (1.42-4.61) * | 1.48(0.66-3.28) |
|  | No | 167 | 112 | 1 | 1 |
| Back pain over the past 30 days | Yes | 17 | 26 | 2.144 (1.12-4.12) * | 0.9(0.35-2.30) |
|  | No | 171 | 122 | 1 | 1 |
| Fever over the past 30 days | Yes | 16 | 32 | 2.966 (1.56-5.65) * | 2.34(0.99-5.53) |
|  | No | 172 | 116 | 1 | 1 |
| Chronic Medical Illness | Yes | 8 | 27 | 5.02 (2.21-11.42) * | 3.32(1.26-8.75) * |
|  | No | 180 | 121 | 1 | 1 |

\* p < 0.05; \*\* P<0.01; 1= Reference; COR= crude odds ratio & AOR= adjusted odds ratio

The odds of having depression on those prisoners who have a history of suicidal attempt is higher than their counterparts (AOR=2.76 CI: 1.04-7.31), and similarly, the odds of having depression is higher on prisoners who sustained a severe stressful life event in the past (AOR=2.57 CI:1.41-4.67). Prisoners with a sentence of 5 to 10 years are more likely to have depression than those sentenced for less than five years (AOR=2.51 CI: 1.32, 4.79) and those who have chronic medical illness also have higher odds to suffer from depression (AOR=2.51 CI:1.32,4.79). (Table 3)

## Discussion

### Prevalence of depression

In this study the prevalence of depression is 44.05% (95% CL: 38.8%, 49.9%), and this finding is consistent with studies done in Ethiopia, Jimma Prison 41.9% and Bahir Dar prison 45.5%; both studies used the same assessment tool (PHQ-9) and similar study set up like that of our research. The result of our study shows a higher prevalence of depression (44.05%) than depression prevalence in general public (4.7% [6] 6.8% -11.0%[39]).

Our study also showed a higher prevalence of depression as compared with studies from countries like Brazil 17.6% [14] and Iran 29%[13], the possible reason might be we used PHQ-9, and both studies used different scale to determine the level of depression, for instance, the study from Iran used DSM-IV Axis and the study from Brazil used MINSI. Besides, unique challenges like inadequate prison facilities that face low-income countries might be the reason for the high prevalence of depression among prisoners in Ethiopia. Studies from South Africa 24.4% [16] and Nigeria 30.8% [18] also revealed a lower level of depression among inmates, the difference could explain this in the way of treatment under prison rule and regulation, and both countries are better in the level of economic development than Ethiopia.

The result of this study is significantly lower than the study done in Egyptian prison in which prevalence’s of depression was 82.5%, this difference may be due to the different instrument used i.e., they used SCL-90 but we used PHQ-9, and the Egyptian study used a sample of 80 participants with convenient sampling [17]. In comparison to our study, a bit higher prevalence of depression, 56.4% was revealed by a study done at Hawassa Central Correctional Institution. This can be due to longitudinal time frame difference of the study period [19].

### Predictors of Depression

The odds of having depression is found to be higher in widowed than in singles (AOR=6.30 CI: 1.09-36.67), this may be due to higher expected parental responsibility and loss of loved ones in widowed. This finding is consistent with the study done in Northwest Amhara, Ethiopia, and a study from Rio de Janeiro, Brazil. [22, 40].

Those prisoners educated at the college level or university level are more likely to have depression than illiterates (AOR= 5.34 CI: 1.59-17.94). Though there are no studies supporting this finding, in our study around 70% of prisoners are attended only primary education and very few proportions (6.3%) of the inmates are attending college or university level education and educated while being a minority in prison may expect better treatment and better respect in the prison while prisons have same treatment irrespective of literacy.

Inline, the study of Northwest Amhara, Ethiopia, the odds of having depression on those prisoners who have a past history of suicidal attempt is higher than their counterparts (AOR=2.76 CI: 1.04-7.31) [22] and suicide is very common phenome with depression and can be part of the depression [7, 41]. Similarly, the odds of having depression is higher on prisoners who sustained a severe stressful life event in the past (AOR=2.57 CI:1.41-4.67), and this finding is consistent with the result from Malesia [42] and a systematic review [43].

As evidenced in different studies, chronic medical illness is highly linked with depression resulting in poor prognosis and suicide [44–46]. Our study also revealed that those prisoners who have chronic medical illness have a higher odd to suffer from depression (AOR=2.51 CI:1.32,4.79) and this is in line with studies from Hawassa and Jimma prisons of Ethiopia [19, 21].

Prisoners with a sentence of 5 to 10 years are more likely to have depression than those sentenced for less than five years (AOR=2.51 CI: 1.32, 4.79) and this finding is supported by a study from Bahir Dar prison (AOR=2.13 CI: 1.01, 5.25). This implies prisoners staying longer in prison are likely to develop depression compared with a short stay at the prison, and this may alarm due mental health attention for those prisoners staying longer than five years in prison.

## Conclusion

This study aimed to measure the prevalence of depression among prisoner of Debre Berhan town prison and to assess factor associated with depression. The findings of our study reported that depression high prevalence of depression among prisoners of Debre Berhan town prison, which accounts for 44.05% of study participants.

This cross-sectional study found that prisoners who are; widowed, educated at college or university level, and prisoners who have a history of suicidal attempt were significantly associated with depression. Furthermore, facing a severe stressful life event, having a chronic medical illness and serving five to ten years of sentence are independently associated with depression.

Therefore, Debre Berhan town prison administration, responsible governmental and non-governmental organizations need to consider early screening and treatment strategies for prisoners who serve a sentence in prison greater than 5years, for those prisoners with a history of suicidal attempt and chronic medical illness to prevent suicide and another adverse effect of depression.

### Limitations of the study

The major limitation of the study was the fact that it was not a multi-centered study. A multi-site study would provide an enormous wealth of information on the prevalence rate of depressive disorder amongst the different prison population in Ethiopia & would enable comparative analysis between them. Another major limitation is that only six females were chosen for this study on random proportional demographic representation, future research with a higher number of female participants might solve the weakness that we faced comparing results by sex.

## Acknowledgment

We want to extend our special gratitude to Debre Berhan University College of Medicine and Health Science for supporting the research project. Our sincere thanks go to Debre Berhan prison officials for their dedicated collaboration and, we are grateful and appreciate all research participants for their unreserved participation.

## Abbreviations

BDI: Beck Depression Inventory
DD: Depressive Disorder
DSM IV-TR: Diagnostic and Statically Manual Four Text Revision
DSM-5: Diagnostic and Statistical Manual of Mental Disorders, Fifth Edition
MDD: Major Depressive Disorder
MINSI: Mini International Neuropsychiatric Interview
PHQ-9: Patient Health Questionnaire- 9
SCI: Structured Clinical Interview
SCL: Psychopathy Symptoms Checklist
SCL-90: Psychiatric Symptoms Checklist-90
SPSS: Statistical Package for Social sciences
WHO: World Health Organization

